# Maternal obesity alters uterine NK cell activity through a functional KIR2DL1/S1 imbalance

**DOI:** 10.1101/167213

**Authors:** Barbara Castellana, Sofie Perdu, Yoona Kim, Kathy Chan, Jawairia Atif, Megan Marziali, Alexander G. Beristain

**Affiliations:** British Columbia Children’s Hospital Research Institute, Vancouver, Canada.; Department of Obstetrics & Gynecology, The University of British Columbia, Vancouver, Canada.

**Author notes:** To whom correspondence should be addressed: Alexander G. Beristain, British Columbia Children’s Hospital Research Institute, Department of Obstetrics & Gynecology, The University of British Columbia, Vancouver, British Columbia, Canada. V5Z 4H4. **CONFLICT OF INTEREST:** The authors declare no conflict of interest.

**Keywords:** Pregnancy, natural killer cell, uterus, decidua, maternal obesity, killer cell immunoglobulin-like receptor

## Abstract

In pregnancy, uterine natural killer cells (uNK) play essential roles in coordinating uterine angiogenesis, blood vessel remodeling, and promoting maternal tolerance to fetal tissue. Deviances from a normal uterine microenvironment are thought to modify uNK function(s), limiting their ability to establish a healthy pregnancy. While maternal obesity has become a major health concern due to associations with adverse effects on fetal and maternal health, our understanding into how obesity contributes to poor pregnancy disorders is essentially unknown. Given the importance of uNK in pregnancy, this study sets out to examine if obesity affects uNK function. Using a cohort of pregnant women, we show that baseline activity of uNK from obese women is elevated, but that enhanced activity does not equate to increased killing potential. Instead, obesity associates with altered uNK production of angiogenic VEGF-A and PlGF. These changes coincide with alterations in NKp46^+^ and NKG2A^+^ uNK subsets and elevated expression of KIR2D(L1/S1/S3/S5) receptors. Detailed examination revealed that obesity leads to imbalances in KIR2DL1/S1 expression that together instruct altered responses to HLA-C2 antigen, including increased production of TNFα. Together, these findings suggest that maternal obesity modulates uNK function by altering angiokine/cytokine production and the response to HLA-C2 antigen.

## INTRODUCTION

Establishment of the fetal-maternal interface in early pregnancy requires coordinated interactions between fetal and maternal cells that help direct placentation and ensure an adequate blood supply to the developing fetus. Early stages of uterine angiogenesis, arterial remodeling, trophoblast invasion & survival, as well as maternal tolerance toward the fetal semi-allograft are controlled in part by maternally-derived uterine immune cells^1,2^. In particular, uterine natural killer cells (uNK), identifiable as CD56^bright^/CD16^−^ cells in humans, constitute the most abundant leukocyte population within the uterus accounting for ~70% of immune cells^2,3^. Unlike peripheral blood NK cells (pbNK), uNK do not normally mount cytotoxic responses, but instead help coordinate placental development by controlling uterine neo-angiogenesis and spiral artery remodeling, as well as limiting the immune response against fetal antigen^4^.

NK activity (i.e. cytotoxicity, cytokine & angiokine production) is controlled in part by activating and inhibiting natural killer receptors (NKRs). NKRs bind epitopes expressed on major histocompatibility (MHC) class I and class I-like molecules, stress-related molecules and cytokines^5–7^. Generally, signaling via activating receptors stimulate cytolytic programs whereas signaling via inhibitory receptors drives immuno-tolerance through mechanisms that dampen activating NKR signal transduction pathways^8^. uNK are phenotypically distinct from pbNK (pbNK are defined phenotypically as CD56^dim^/CD16^+^) in that they express a distinct repertoire of NKRs that facilitate interaction with trophoblast-derived MHC class I molecules (HLA-C, HLA-E, and HLA-G) but not HLA-A or HLA-B^9^. Depending on the activating/inhibitory NKR balance, interaction of uNK with MHC ligand can promote or restrain uNK activity. It is suggested that inadequate (not enough) or inappropriate (too much) uNK activation contributes to preterm birth, recurrent miscarriage, and preeclampsia by limiting angiogenesis/artery remodeling or over-sensitizing uNK to being cytotoxic^10–16^.

Obesity, with a prevalence between 13-18% in women of reproductive age^17^, correlates with higher incidences of poor pregnancy outcomes^18,19^. For example, pre-existing obesity associates with recurrent miscarriage^20^, gestational diabetes^21^, preterm birth^22^, and preeclampsia^23^. Studies designed to generate insight into how maternal obesity contributes to poor pregnancy outcomes have shown in rodents that high-fat diet exposure prior to and during pregnancy results in impaired uterine artery remodeling^24,25^. Notably, vascular defects of this type are also observed in rats and mice deficient in uNK^26^. Consistent with these observations, previous findings from our group show that maternal obesity in women associates with impaired uterine artery remodeling^27^. Importantly, we show that vascular changes coincide with fewer uNK and altered global uNK gene expression^27^. Together, these studies suggest that obesity may modify pro-angiogenic and vascular-remodeling programs controlled by uNK within the maternal-fetal interface. However, little is known about how maternal obesity affects uNK biology.

In this study we examine if maternal obesity alters uNK activity and function. We show that uNK from obese women are more active and secrete different amounts of the angiogenic factors PlGF and VEGF-A. Phenotypic analyses show altered expression of NKRs in obese women compared to lean women, where differences in activating (NKp46, KIR2DS1) and inhibitory (NKG2A, KIR2DL1) NKR cell populations are observed. Importantly, we provide evidence that the increase in obesity-linked uNK activity is in part due to changes in expression of the NKRs KIR2DL1 and KIR2DS1. We show that upon HLA-C2 stimulation, KIR2DS1-expressing uNK from lean and obese women differentially regulate TNFα production. Together, this work provides insight into how maternal obesity affects uNK function. Further, this work identifies uNK-directed processes modified in obesity that may contribute to impaired vascular remodeling, placental function and pregnancy outcome.

## RESULTS

### Maternal obesity promotes uNK activation

Our lab previously reported that maternal obesity associates with reductions in uNK cell numbers and coincides with impaired uterine blood vessel remodeling^27^. To examine if these changes might be linked with altered uNK activity, we compared degranulation/activation responses in uNK isolated from lean (BMI 20-24.9 kg m^−2^) and obese (BMI ≥30 kg m^−2^) women in their first trimester of pregnancy (Table 1). Following culture in the presence or absence of phorbol 12-myristate 13-acetate/ionomycin (PMA), uNK activity was measured via surface expression of the degranulation marker CD107a^28^. For these studies, uNK were defined as CD56^bright^/CD9^+^/CD16^−^/CD3^−^ CD45^+^ cells (Figure 1a). As expected, PMA stimulation resulted in potent uNK activation in both BMI groups (Figure 1b,c). However, proportions of CD107a^+^ uNK from obese women were more abundant in both unstimulated (median = 5.4*% vs* 2.6%) and stimulated (median = 11.8% *vs* 5.9%) conditions (Figure 1c). These initial findings indicate that maternal obesity associates with a heightened state of uNK activation.

**Figure 1.**
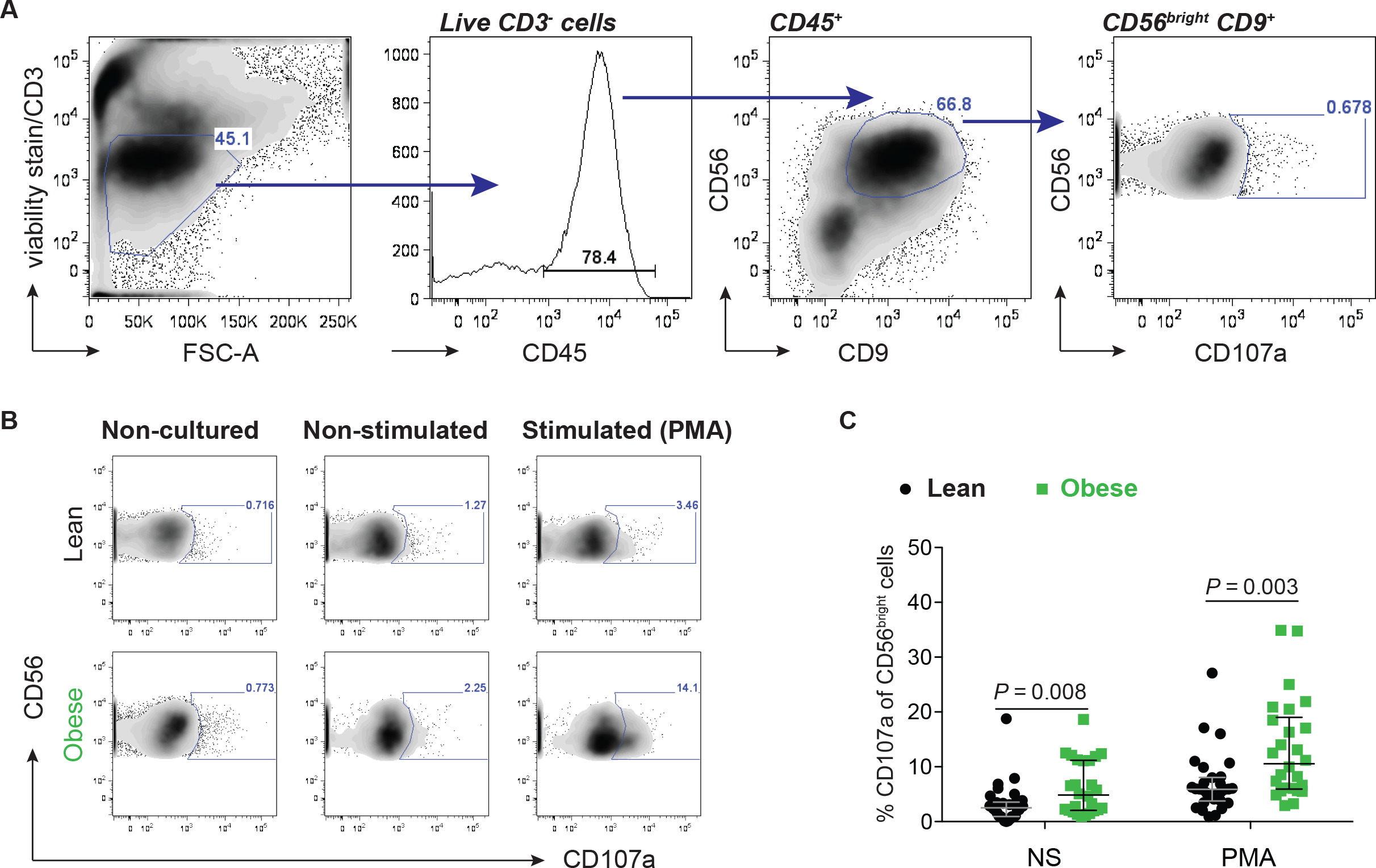
Maternal obesity associates with heightened uNK activity. **(a)** Flow cytometry gating strategy used to analyze degranulation in uNK defined as CD56^bright^/CD9^+^ cells. **(b)** Representative flow cytometry plots of CD107a in uNK cells at baseline (non-cultured), or cultured in the absence (non-stimulated) or presence (stimulated) of PMA/ionomycin for 4 hours. Baseline measurements were determined from *ex vivo* non-cultured uNK. Percentages of CD56^bright^/CD9^+^ cells positive for CD107a are indicated within the plots. **(c)** Scatter plots depicting percentages of CD56^bright^/CD9^+^ uNK from lean (black circles; n = 30) and obese (green squares; n = 26) expressing CD107a following PMA/ionomycin treatment. *P* values (nonparametric two-tailed Mann-Whitney *t* test) are shown.

**Table 1.** Clinical characteristics

|  | Average<br>age | IQR Age | Average<br>GA | IQR GA | Average<br>BMI | IQR BMI |
| --- | --- | --- | --- | --- | --- | --- |
| Lean women<br>(N=104) | 27.84 | 23 to 33 | 7.7 | 6.5 to 8.5 | 22.05 | 20.79 to 23.49 |
| Obese women<br>(N=80) | 28.36 | 23 to 33<br>*P* = 0.32 | 8.5 | 7.1 to 9.6<br>*P* = 0.003 | 34.56 | 31.62 to 37.04<br>*P* < 0.0001 |

### Obesity associates with altered uNK angiokine production

To gain insight into how obesity-linked differences in uNK activity translate into uNK functional differences, we measured by flow cytometry the intracellular production of tumor necrosis factor α (TNFα) and interferon gamma (IFNγ), two cytokines expressed by uNK known to play central roles in uterine blood vessel remodeling^29^. In line with higher CD107a^+^ uNK proportions, a non-significant but trending increase in baseline TNFα production was observed in uNK from obese women (*P* = 0.05); baseline IFNγ levels between BMI groups were similar (Figure 2a-c). Following PMA stimulation, uNK from both groups markedly enhanced production of TNFα and IFNγ (Figure 2c). Although we did not detect a significant difference in cytokine expression between BMI groups, uNK from obese women again showed a slight tendency to produce more TNFα (median = 19.4% *vs* 9.9%, *P =* 0.05) (Figure 2c).

**Figure 2.**
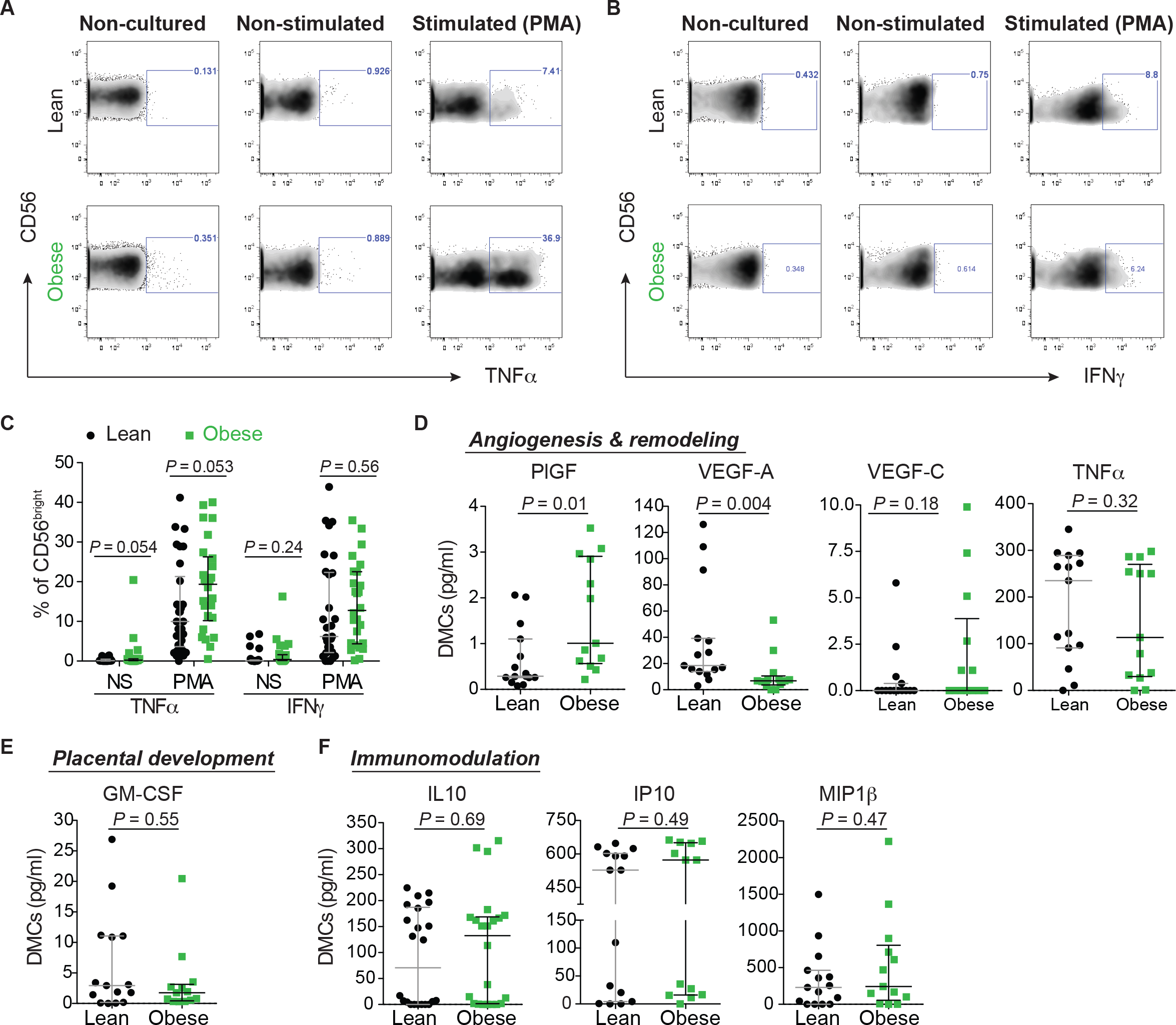
Maternal obesity leads to alterations in uNK cytokine production. Representative flow cytometry plots of **(a)** TNFα and **(b)** IFNγ expression in uNK from lean (n = 30) and obese (n = 26) women at baseline (non-cultured) or cultured in the absence (non-stimulated) or presence (stimulated) of PMA/ionomycin for 4 hours. Percentages of CD56^bright^ cells positive for TNFα and IFNγ are indicated within plots. **(c)** Scatter plots showing median and IQR values of proportions of CD56^bright^ uNK expressing TNFα and IFNγ from lean (black circles) and obese (green squares) women. Scatter plots depicting levels of secreted **(d)** angiogenic and remodeling [PlGF, VEGF-A, VEGF-C, TNFα], **(e)** placental development [GM-CSF], and **(f)** immunomodulation [IL10, IP10, MIP1β] factors from decidual leukocytes (>60% uNK) derived from lean (black; n = 15) and obese (green; n = 13) women. *P* values (nonparametric two-tailed Mann-Whitney *t* test) are shown.

To complement the above intracellular cytokine measurements, we also measured the secretion of 8 factors from conditioned media (CM) generated by uNK-enriched *ex vivo* cultures (> 60% uNK) from lean and obese women. These factors included: granulocyte-macrophage colony-stimulating factor (GM-CSF), interleukin 10 (IL10), interferon-γ inducible protein 10 (IP10), macrophage inflammatory protein 1β (MIP1β), placental growth factor (PlGF), TNFα, and vascular endothelial growth factor A and C (VEGF-A, VEGF-C). These cytokines were chosen based on prior studies showing their production by uNK and their importance in controlling pregnancy-related processes^30,31^. uNK CM from obese women secreted lower amounts of VEGF-A (median = 6.8 pg/ml *vs* 18.4 pg/ml) and higher amounts PlGF (median = 1.0 pg/ml *vs* 0.3 pg/ml) (Figure 2d). However, secretion of TNFα, VEGF-C, GM-CSF, IL10, IP10 and MIP1β were comparable between uNK CM from either BMI group (Figure 2d-f). Our results suggest that obesity-linked changes in uNK activity translate into differences in angiokine secretion, reflecting possible modifications to angiogenic processes.

### Maternal obesity associates with phenotypical differences in NKR expression

Our finding that maternal obesity associates with heightened uNK activity and altered angiokine production indicates that differences in NKR expression/abundance may exist. To examine this, we compared the expression of 8 NKRs in uNKs from lean and obese women using a flow cytometry approach outlined in Supplementary Figure 1a. Specifically, we measured activating (NKG2D) and inhibitory (NKG2A) CD94/NKG2 family members, natural cytotoxicity receptors NKp30, NKp44 and NKp46, inhibitory leukocyte immunoglobulin-like receptor (LILRB) 1, and members of the killer-cell immunoglobulin-like receptor (KIR)2D subfamily [using antibodies targeting KIR2DL4 and KIR2D(L1/S1/S3/S5)]. These NKRs were selected in part due to their importance in controlling NK cell activation^5,6,8^, their prior description in uNK^30,32^, or their association with pregnancy-related outcome^16,33,34^. uNK from lean women ubiquitously expressed inhibitory NKG2A and activating NKp30 and NKp46, while varied proportions of NKG2D^+^ (58.9%), LILRB1^+^ (30.1%), and KIR2DL4^+^ (62.0%) uNK were observed (Figure 3a-d, and S1b-e). Between BMI groups, proportions of NKG2D^+^, NKp30^+^, NKp44^+^, LILRB1^+^, and KIR2DL4^+^ uNK were comparable, however, maternal obesity associated with a decrease in NKG2A^+^ (93.3% *vs* 96.6%) and NKp46^+^ (68.3% *vs* 96.0%) cell proportions (Figure 3a,b). Interestingly, the decrease in NKp46^+^ cell proportions in obese women was characterized by a distinct bimodal distribution (Figure 3b). Additionally, uNK from obese women had elevated proportions of KIR2D^+^ [L1/S1/S3/S5; herein referred to as KIR^+^] uNK (46.0% *vs* 35.6%) (Figure 3c). These differences in NKR proportions in obese women also associated with decreases in cell surface levels (median fluorescence intensity; MFI) of NKp46 and NKG2A (NKp46: 1529 arbitrary units (AU) *vs* 2567 AU; NKG2A: 1349 AU *vs* 2051 AU) (Figure 3a,b). These results suggest that uNK from obese women show a distinct NKR phenotype defined by changes in both activating and inhibitory NKRs.

**Figure 3.**
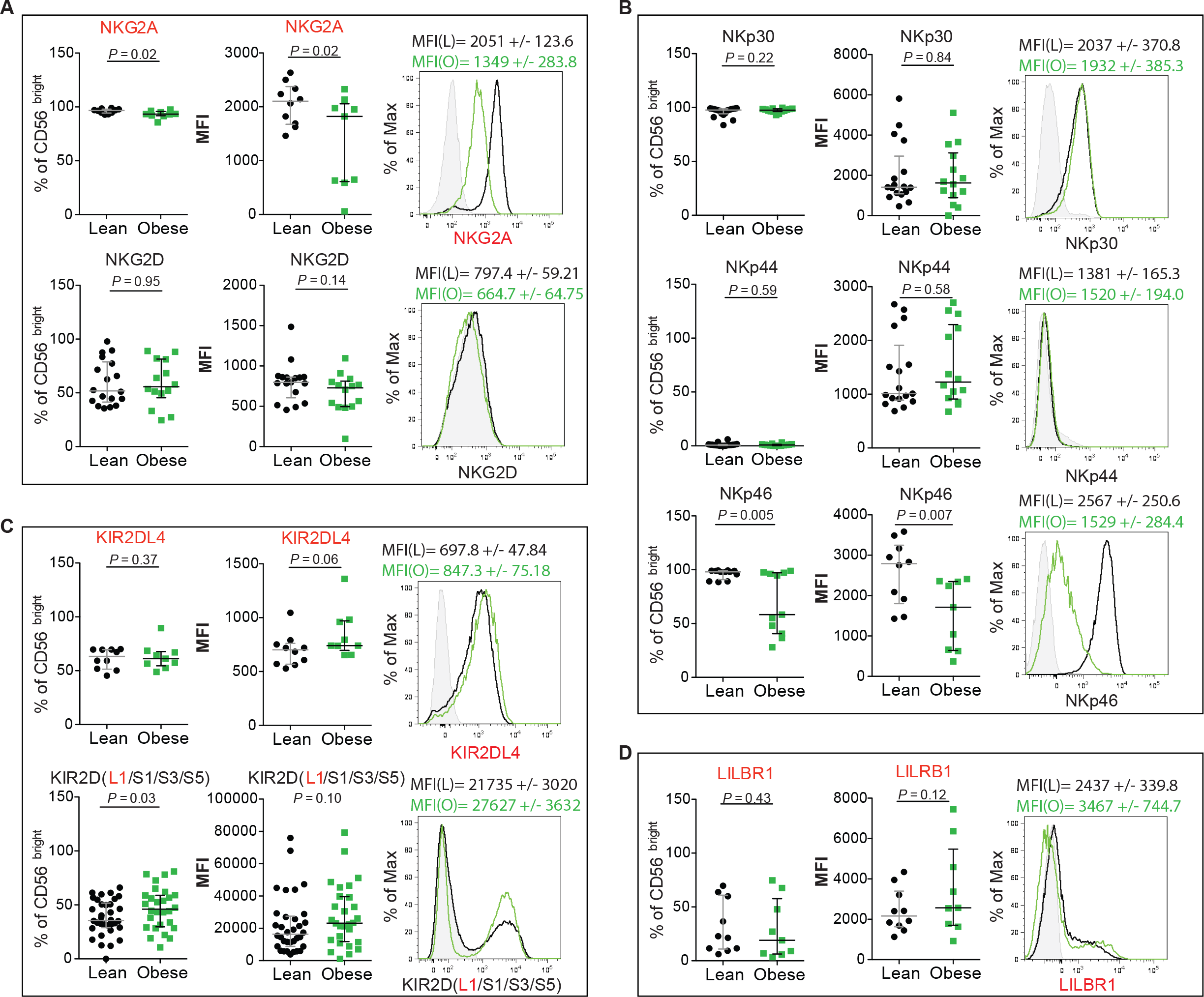
Maternal obesity correlates with changes in activating and inhibitory Natural Killer Receptors (NKRs). Scatter plots show flow cytometry-derived proportions of uNK (gated on CD56^bright^ cells) expressing **(a)** inhibitory NKG2A (n = 9 lean *vs* 10 obese) and activating NKG2D (n =14 lean *vs* 18 obese) CD94/NKG2 family receptors, **(b)** activating natural cytotoxicity receptors NKp30 (n = 14 lean *vs* 18 obese), NKp44 (n = 14 lean *vs* 18 obese), and NKp46 (n = 11 lean *vs* 12 obese), **(c)** inhibitory KIR2DL4 (n = 9 lean *vs* 10 obese) and KIR2DS1/S3/S5 (n = 28 lean *vs* 34 obese) killer-cell immunoglobulin-like receptor (KIR)2D subfamily receptors, and **(d)** the leukocyte immunoglobulin-like receptor LILRB1 (n = 9 lean *vs* 10 obese), as well as their surface expression levels (MFIs). On the right of each scatter plot, representative histograms display uNK surface levels of individual NKRs. Grey area indicates unstained control, where black and green histograms indicate representative lean and obese subject expression levels. MFIs and standard deviation (SD) of lean (L) and obese (O) samples are shown above the histogram. Activating (black) and inhibitory (red) receptors are color-coded. *P* values (nonparametric two-tailed Mann-Whitney *t* test) are shown.

### Maternal obesity drives KIR2DL1/S1 NKR imbalances

KIRs are involved in NK cell education, promotion of fetal tolerance and contribution to successful placentation^35,36^. Because previous studies have highlighted strong associations with pregnancy outcome and KIR2DL1/S1 expression^35^, and because we showed that maternal obesity associates with increased proportions of KIR2D^+^ uNK, we next set out to examine if maternal obesity leads to imbalances in inhibitory KIR2DL1 (2DL1) or activating KIR2DS1 (2DS1). Using a flow cytometry approach that identifies distinct 2DL1 and 2DS1 single-positive (sp) and mixed double-positive (dp) uNK populations (Figure 4a), we measured proportions and surface expression levels of these KIRs in lean and obese women. Analysis by qPCR revealed that maternal obesity correlates with higher mRNA levels of *KIR2DS1* in uNK (purified by negative selection), while levels of inhibitory *KIR2DL1* and activating *KIR2DS3/S5* did not differ between BMI groups (Supplementary Figure S2).

**Figure 4.**
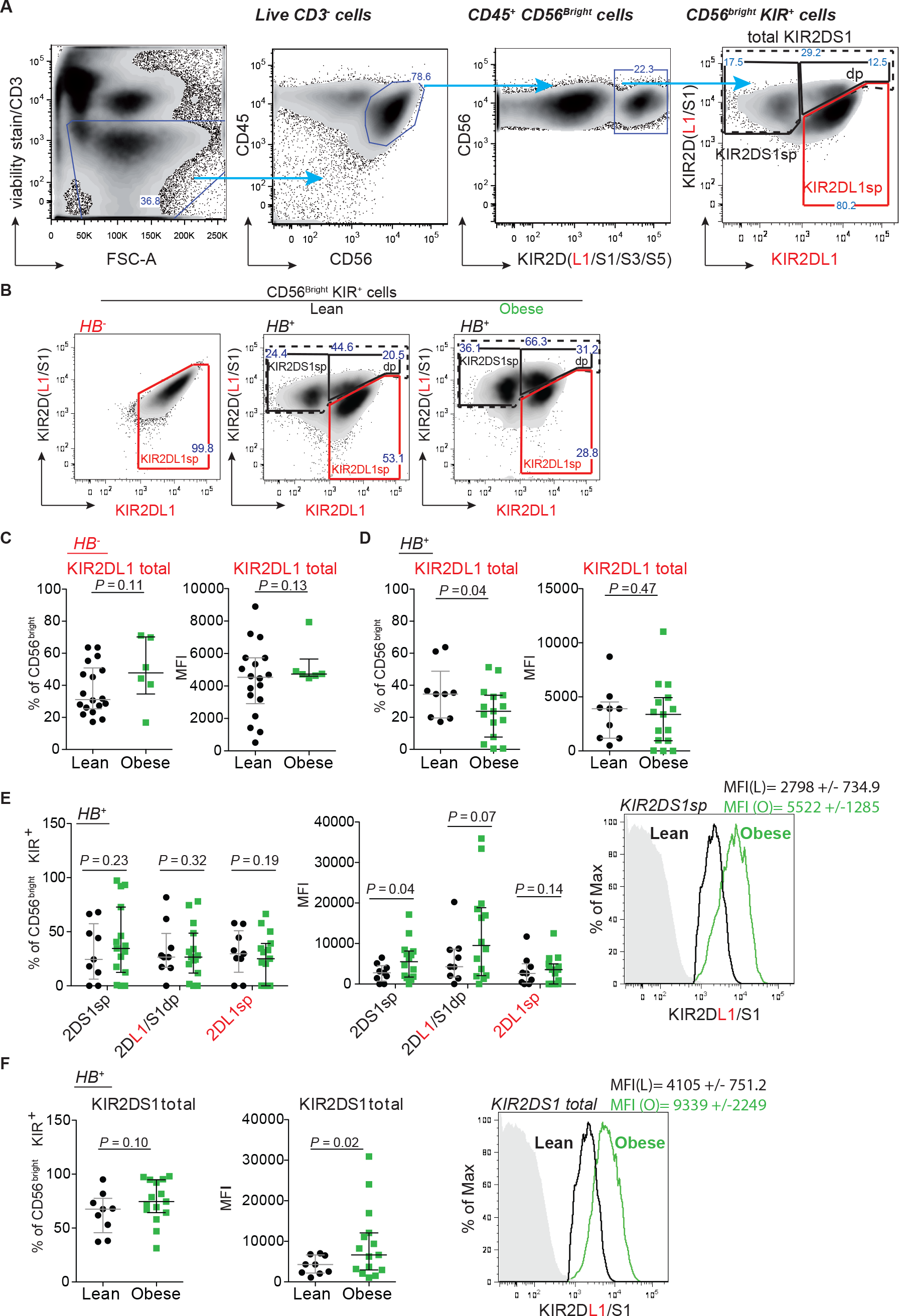
Obesity links with imbalances in uNK KIR2DS1/L1 expression. **(a)** Shown is the flow cytometry gating strategy for KIR2DL1/S1 analysis. CD3/FVD780 exclusion identifies live CD3^−^ cells that are further selected on CD45, CD56, and KIR2D(L1/S1/S3/S5) positivity. Within the KIR^+^ population, uNK are analyzed for KIR2DS1 and KIR2DL1 expression using antibodies directed against KIR2DL1 and KIR2DS1/L1 where three subsets (KIR2DS1sp, KIR2DL1/S1dp, KIR2DL1sp; sp, single-positive; dp, double-positive) are identified. **(b)** Representative flow cytometry plots of KIR2DS1 and KIR2DL1 uNK populations from a KIR2DS1^−^ haplotype B negative subject (HB^−^; left) and KIR2DS1^+^ haplotype B positive (HB^+^) subjects from lean (n = 9) and obese (n = 15) women. The percentage of cells is shown in each gated area. **(c-d)** Scatter plots show total KIR2DL1 proportions of uNK (gated on CD56^bright^ cells) and expression levels (MFI) from HB^−^ and HB^+^ lean and obese women. **(e)** Scatter plots show KIR2DS1 proportions (2DS1sp, 2DL1/S1dp, and 2DL1sp) and surface levels (MFI) in uNK (gated on CD56^bright^/KIR^+^) in HB^+^ lean and obese women. Representative flow cytometry histogram (right) shows KIR2DS1 levels (MFI) in KIR2DS1sp uNK. **(f)** Percentage and expression of total KIR2DS1 (combined KIR2DS1sp and KIR2DL1/S1dp populations) in HB^+^ lean and obese subjects. MFI and standard deviation (SD) of KIR2DS1 is shown: solid grey area indicates the fluorescence minus one (FMO) baseline signal; black and green histograms indicate 2DS1 MFI in lean and green subjects. *P* values (nonparametric two-tailed Mann-Whitney *t* test) are shown.

As the presence or absence of 2DS1 depends on inherent maternal and paternal haplotype combinations [two main haplotypes exists: haplotype A (HA), primarily comprised of inhibitory receptors (i.e. lacking 2DS1), and haplotype B (HB), containing both activating and inhibitory receptors, including 2DS1^37^], we sub-stratified lean and obese cohorts into KIR HB^−^ (2DL1^+^/S1^−^) and HB^+^ (2DL1^+^/S1^+^) by PCR genotyping (Supplemental Table 1). Flow cytometry analysis revealed that HB^−^ uNK are negative for 2DS1, whereas mixed 2DL1^+^/2DS1^+^ populations are seen within HB^+^ cells in both lean and obese women (Figure 4a,b). Among HB^−^ individuals (> 95% are allotypically 2DL1^38^), total proportions of 2DL1^+^ uNK were similar between lean and obese women (Figure 4c). Interestingly, uNK from obese HB^+^ women showed reduced proportions of 2DL1^+^ cells (gated on total CD56^bright^ uNK; median = 23.8% *vs* 34.5%). In either haplotype, 2DL1 MFI did not differ between lean and obese groups (Figure 4d).

Focusing on only HB^+^ (containing *2DS1* gene) uNK, we again examined 2DL1/S1 subsets within KIR^+^ cells in both BMI groups. Proportions of 2DS1sp and dp populations did not differ between lean and obese groups (Figure 4e). However, 2DS1 surface expression (MFI) within the 2DS1sp population in obese women was approximately 2-fold higher; levels of 2DL1 within 2DL1/S1dp and 2DL1sp populations did not differ (Figure 4e). Measuring 2DS1 MFI on all KIR^+^ uNK (combined 2DS1sp and 2DL1/S1dp populations) showed that, similar to expression within the 2DS1sp population, 2DS1 levels are elevated in uNK from obese women (Figure 4f). Together our results indicate that maternal obesity instructs changes in KIR2D expression to preferentially favour 2DS1, potentially conferring functional changes towards increased cell activation.

### KIR2DS1^+^ uNK from obese women show enhanced activity

Because KIR expression is influenced by maternal KIR haplotype, we next revisited our findings (Figure 1) that examined if differences in uNK activity exist between lean and obese women by stratifying women into HB^−^ and HB^+^ haplotypes. In non-stimulated conditions, the proportion of CD107a^+^ HB^−^ uNK was not different between BMI groups [1.8% (obese) *vs* 2.1% (lean)]. However, within HB^+^ uNK, maternal obesity associated with higher proportions of CD107a^+^ cells (6.7% *vs* 2.7%; Figure 5a). Following PMA stimulation, uNK from obese women of either haplotype degranulated at higher frequencies (HB^−^: 8.2% *vs* 5.2%; HB^+^: 17.1% *vs* 8.2%), where HB^+^ uNK cells from obese women showed modestly higher degranulation rates than HB^−^ cells from obese women (Figure 5b). Surprisingly, proportions of TNFα-expressing uNK were higher in PMA-primed HB^−^ cells from obese women, where BMI-associated differences were not seen in HB^+^ cells (Figure 5c). Consistent with our previous results, maternal obesity did not affect proportions of IFNγ expressing uNK, regardless of KIR haplotype (Figure 5c).

**Figure 5.**
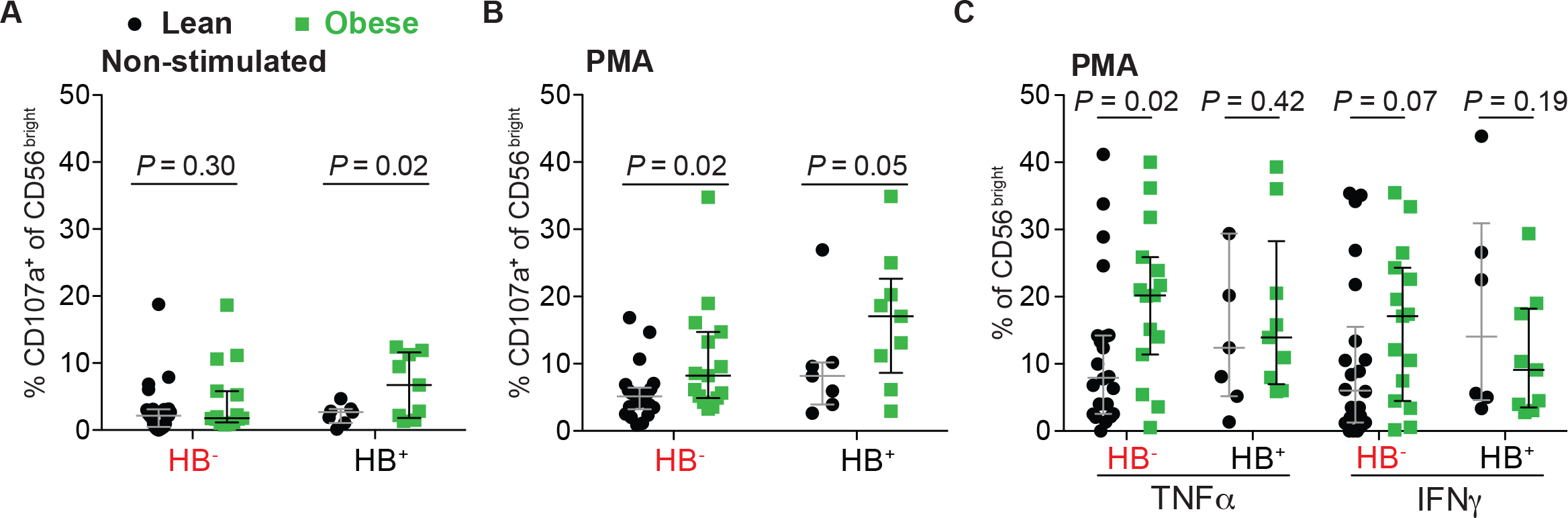
Maternal obesity and KIR haplotype interact to potentiate uNK activity. **(a)** Scatter plots show flow cytometry-derived proportions of CD107a cells in **(a)** non-stimulated and **(b)** PMA-stimulated uNK cultures from HB^−/+^ lean (black: HB^−^ n = 22; HB^+^ n = 7) and obese (green: HB^−^ n = 15; HB^+^ n = 9) subjects. **(c)** Scatter plots show proportions of uNK from HB^−/+^ lean and obese subjects producing TNFα and IFNγ following 4 hours of PMA stimulation. *P* values (nonparametric two-tailed Mann-Whitney *t* test) are shown.

### Obesity, combined with maternal HLA-C2 allotype, potentiates uNK activity

KIR expression can be affected by gestational age^39^ and maternal HLA-C^40^. To determine if KIR changes may relate to these BMI-independent variables, we next examined if gestational age and maternal HLA-C affects 2DL1/S1 uNK proportions or expression levels. uNK 2DL1/S1 frequency and expression level was comparable amongst different gestational ages (Supplementary Figure S3). Homozygous C1/C1 subjects showed an inverse correlation in 2DS1 and 2DL1 frequency depending on BMI. Specifically, uNK cells from obese C1/C1 women exhibited higher proportions of 2DS1^+^ cells and lower proportions of 2DL1^+^ cells (Supplementary Figure 4b,c). Contrary to these KIR imbalances, we also observed that the presence of maternal HLA-C2 associates with an increase in degranulation in non-stimulated uNK from obese women (compared to C1/C1); this association was not maintained following PMA stimulation (Figure 6a,b). The impact of C2 allele on HB^+^ cells was clear after PMA-priming (Figure 6c). However, HLA-C2-associated heightened activity did not translate into an increase in PMA-stimulated cytokine production, although a modest trend was observed (Figure 6d). These results demonstrate that in the context of obesity, maternal HLA-C2 allele equates to heightened activity in HB^+^ uNK.

**Figure 6.**
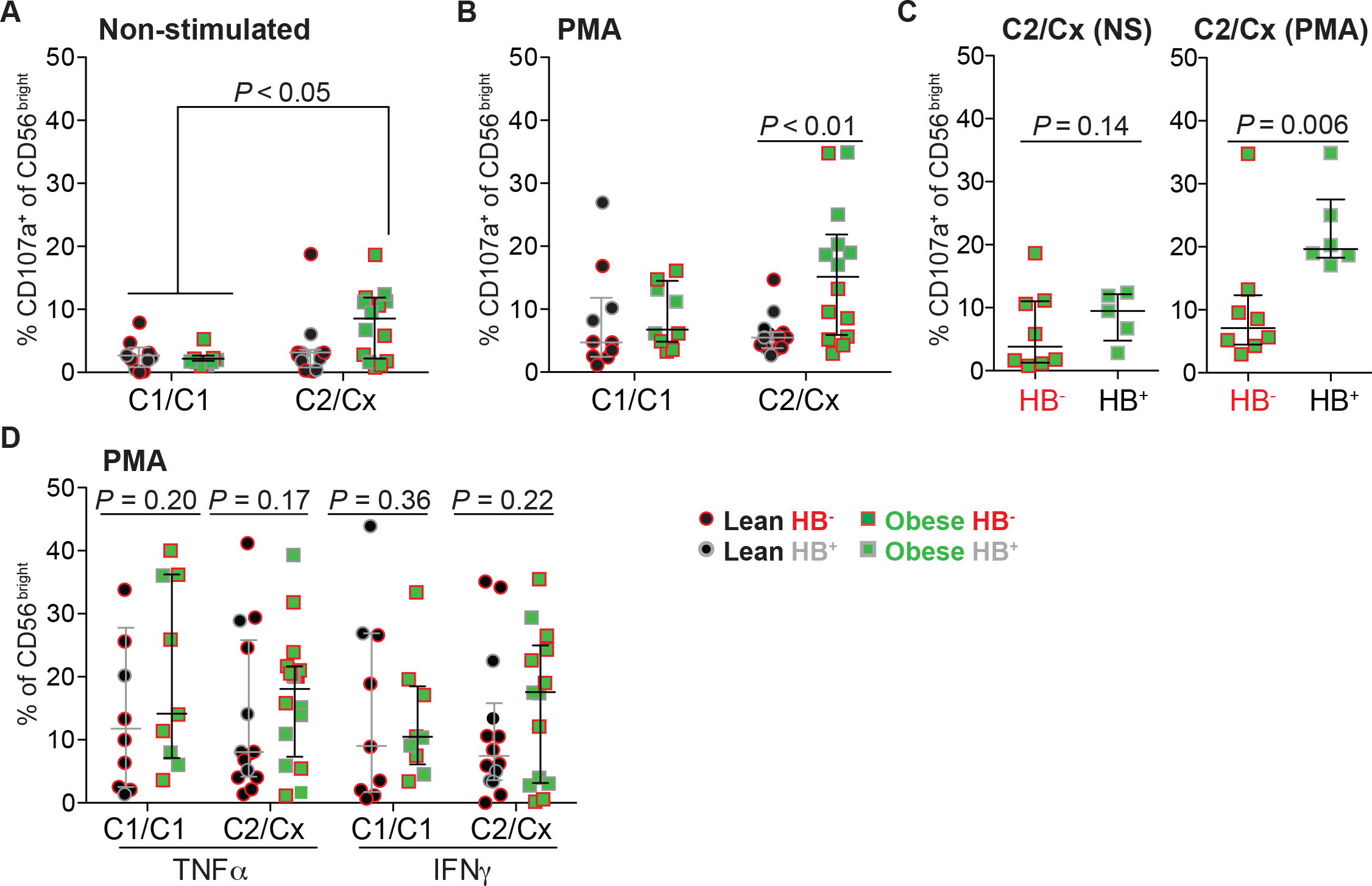
Maternal HLA-C2 and KIR2DS1 correlate with an increase in uNK activity. Scatter plots show proportions of uNK from HB^−/+^ and HLA-C1/C2 genotyped lean (HB^−^ n = 6; HB^+^ n = 8) and obese (HB^−^ n = 8; HB^+^ n = 6) women expressing CD107a following **(a)** no treatment (non-stimulated) and **(b)** PMA/ionomycin (PMA) stimulation. Cx indicates that the individual is either HLA-C2 homozygous or heterozygous. uNK from lean or obese subjects are indicated as black circles or green squares, while HB haplotype are indicated as red outline (HB^−^) or grey outline (HB^+^). **(c)** As above, except scatter plots show comparisons of CD107a^+^ proportions of uNK from only HB^−^ or HB^+^ obese subjects. **(d)** Proportions of TNFα^+^ and IFNγ^+^ uNK cells from HLA-C1/C2 and HB^−/+^ genotyped lean and obese women after PMA treatment. NS = non-stimulated. *P* values (One-way ANOVA-Dunn’s multiple comparisons-[panels **a,b**] test and nonparametric two-tailed Mann-Whitney *t* test [panels **c,d**]) are shown.

### uNK from obese women show enhanced HLA-C2-induced activity

To examine if KIR2DL1/S1 alterations in obese/HB^+^ uNK translate into functional changes, we measured uNK responsiveness towards ectopically expressed HLA-C2 (Supplementary Figure S5a). To this end, HLA class I-deficient K562 cells stably expressing the HLA-Cw*0602 allele were subjected to uNK activity/killing assays using KIR-genotyped uNK from lean and obese groups. HLA-Cw*0602 is the fourth most frequent HLA-C2 allele^41^ and its specificity for 2DL1/S1 allowed us to minimize confounding effects elicited by other HLA-C2-receptors (i.e. LILRB1, KIR2DL2/L3, and KIR2DS4)^42^. To control for KIR-HLA-C2 interaction and to enable differentiation between 2DS1/2DL1-directed responses against HLA-C2, CD107a surface mobilization in CD56^bright^/KIR^−^ cells and HB^−^ uNK were compared to KIR^+^/HB^+^ cells (Figure 7a and S5b). As expected, CD107a mobilization within CD56^bright^/KIR^−^ uNK (HB^−^ and HB^+^) in response to HLA-C2-expressing K562 cells was minimal (Supplementary Figure S5b). Similarly, HB^−^ KIR^+^ uNK (lacking 2DS1) from both BMI groups were also minimally responsive towards HLA-C2 (Figure 7b). Importantly, 2DS1^+^ cells from HB^+^ obese women showed significantly higher CD107a mobilization than 2DS1^+^ cells from lean women in response to high levels of HLA-C2 expression (4.2% *vs* 0.3%). While low-level HLA-C2 expression in target cells showed a trend for increased degranulation in obese 2DS1-expressing uNK, this was not significant. Together, these findings suggest that obesity alters the 2DL1/S1 balance in favour of instructing an activating signal (Figure 7b). Interestingly, HB^+^ uNK from obese C1/C1 women, which have lower levels of 2DL1 and increased proportions of 2DS1 (Supplementary Figure S4b,c), had higher degranulation rates compared to HB^+^ uNK from lean C1/C1 women after HLA-C2 stimulation (Supplementary Figure 5c). This suggests that the absence of a maternal HLA-C2 allele may affect KIR2DS1/L1 responsiveness (education/licensing^43^) to its cognate ligand.

**Figure 7.**
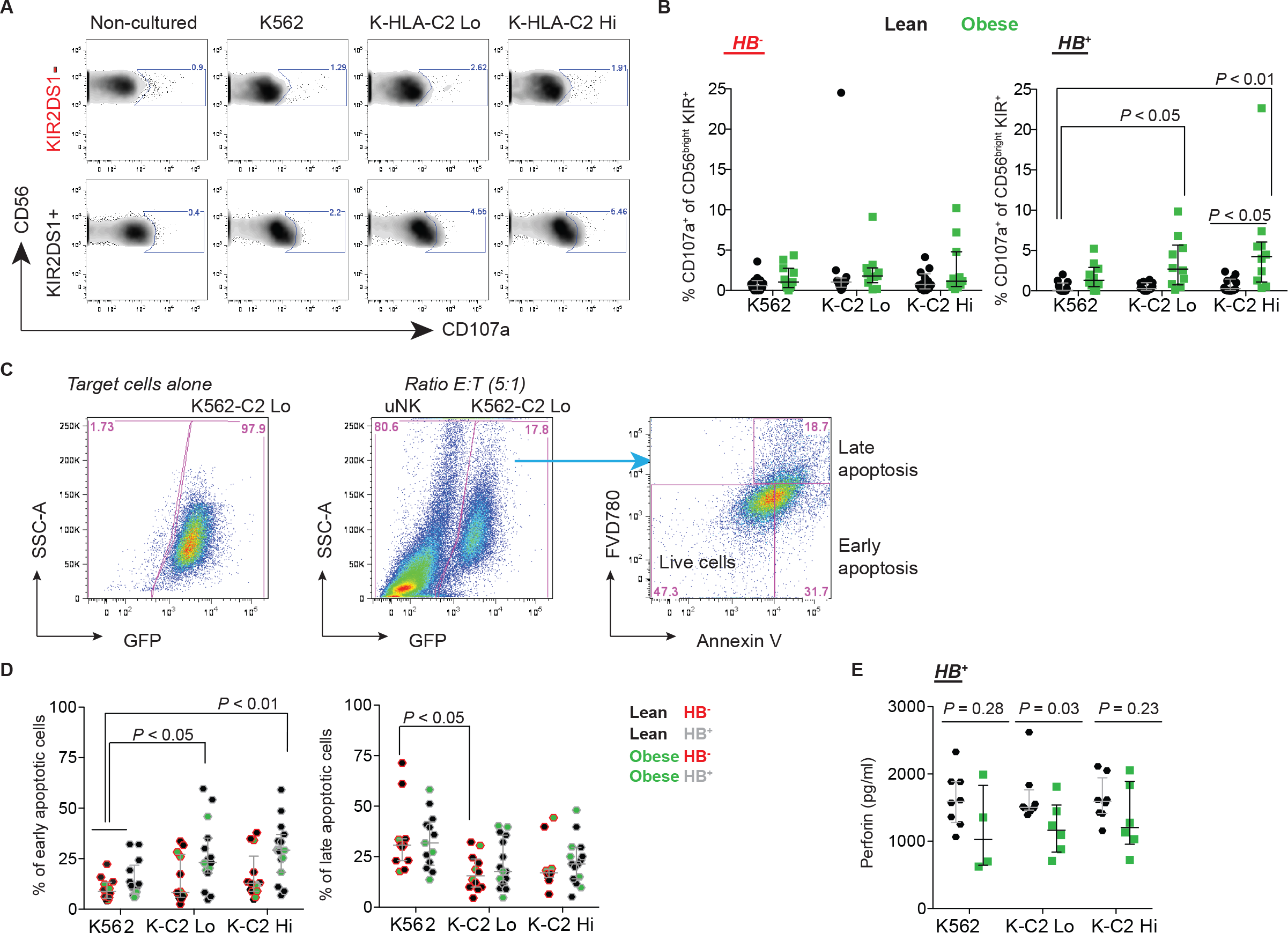
HLA-C2 potentiates heightened activity in KIR2DS1^+^ uNK from obese women. **(a)** Representative flow cytometry plots showing proportions of uNK from lean (HB^−^ n = 13; HB^+^ n= 12) and obese (HB^−^ n= 5; HB^+^ n= 10) women expressing CD107a following 4 hours of co-culture with K562 target cells (5:1 E/T) ectopically expressing HLA-C2-eGFP (HLA-Cw*0602). K562 clones expressing ectopic HLA-C2 at low (K-C2 Lo) or high (K-C2 Hi) levels were established; K562 cells transduced with an empty vector (K562) served as a control. **(b)** Scatter plots show proportions of CD56^bright^/KIR^+^ uNK from HB^−^ (left) and HB^+^ (right) lean and obese women expressing CD107a. **(c)** Shown is the gating strategy for analyzing early and late apoptosis in K562 cells co-cultured with decidual leukocytes (> 60% uNK) for 4 hours. Early and late apoptosis in K562 cells is measured by annexin V and fixable viability dye (FVD) single or dual positivity. **(d)** Scatter plots depict the proportion of early (left) and late (right) apoptotic K562 target cells following co-culture with HB^−^ (red outline; n = 13) or HB^+^ (grey outline; n = 13) uNK from lean (black filled) or obese (green filled) women. **(e)** Scatter plot show levels of secreted perforin from decidual leukocytes from lean and obese women in response to co-culture with HLA-C2-expressing K562 target cells. One-way ANOVA (Dunn’s multiple comparisons test) were performed for panels **(b-d)**. *P* values (nonparametric two-tailed Mann-Whitney *t* test) are shown for panel **(e)**.

Activation of KIR2DS1 strongly triggers cytolysis and cytokine production in pbNK, and this engagement in cytotoxicity is more striking in the absence of KIR2DL1^44–46^. As our above finding shows that HLA-C2 engagement enhances degranulation in HB^+^ uNK from obese women, we next examined whether HLA-C2 interactions also induce cytotoxicity. As expected, HLA-C2 interaction with HB^+^ uNK resulted in increased early apoptosis (measured by annexin V) of K562 target cells compared to culture with HB^−^ uNK cells (K562: 14.9% *vs* 9.8%; K-C2 Lo: 26.5% *vs* 14.2%; K-C2 Hi: 28.9% *vs* 16.0%); however we did not observe differences in late apoptosis based on KIR haplotype (Figure 7c,d). Moreover, within HB^+^ uNK, maternal obesity seemed to associate with impaired HLA-C2 instructed K562 killing, although low sample size/power prevented conclusive interpretation (Figure 7c,d). To complement annexin V killing assays, we also analyzed whether secreted perforin levels were different among lean and obese HB^+^ subjects. In agreement with the annexin V results, the levels of secreted perforin were comparable among BMI groups (Figure 7e). However, obesity appeared to dampen perforin secretion, suggesting that maternal obesity results in impaired uNK cytotoxicity. To exclude the contribution of other KIR-expressing cells in HLA-C2 directed cytolysis, we also analyzed proportions of T (CD3^+^) and NKT (CD56^+^/CD3^+^) cells and their degranulation upon HLA-C2 stimulation. We found the contribution of these cells on cytotoxicity was negligible (Supplementary Figure S6). These findings suggest that although HB^+^ uNK cells from obese women show enhanced HLA-C2 instructed activity (CD107a mobilization), this does not translate into enhanced cytotoxicity.

### Skewing of 2DS1 in lean and obese women results in dichotomous TNFα production

To investigate if the obesity-linked KIR2D-associated increase in uNK activity may also translate into altered production of cytokines/angiokines important in pregnancy, we measured factors from our cytokine 8-plex panel in conditioned media of HB^+^ and HB^−^ uNK from lean and obese women stimulated with a KIR2DL1/S1 crosslinking/activating antibody (clone 11BP6). Changes in cytokine secretion in uNK activated with 11BP6 were compared to responses from isotype-matched IgG1 antibody treatment within paired uNK cultures. As expected, 11BP6 treatment in HB^+^ uNK led to CD107a surface induction, whereas crosslinking did not lead to degranulation responses in HB^−^ cells; these findings indicate that 11BP6 is only capable of inducing an activating response in uNK expressing 2DS1 (Figure 8a). Within lean HB^+^ uNK, 11BP6 crosslinking resulted in an overall decrease in TNFα; two 2DS1sp-containing uNK preparation responded to HLA-C2 by promoting TNFα secretion (Figure 8b). As expected, secretion of TNFα in HB^−^ uNK from either lean or obese groups was not affected (Supplementary Figure 7a). Additionally, we did not observe consistent changes in PlGF, VEGF-A, VEGF-C, GM-CSF, IL10, IP10 or MIP1β following 2DL1/2DS1 crosslinking in uNK from lean HB^+/−^ women (Figure 8b-d and S7). While a trending increase in GM-CSF production was observed in HB^+^ uNK from obese women following crosslinking, penetrant and consistent increases in TNFα production were observed (Figure 8b,d). Notably, a significant interaction between KIR2DL1/S1 crosslinking and BMI was identified, where 11BP6 had opposing effects on TNFα production in uNK from lean and obese women (Figure 8b). Taken together, KIR2DL1/S1 crosslinking in uNK cultures from obese/HB^+^ women promoted TNFα secretion and also appeared to dampen the inhibitory effect of cytokine secretion (i.e. PlGF and TNFα) that was observed in lean HB^+^ cells. These data suggest that the uNK KIR2DL1/S1 imbalance in obese women, potentially favouring 2DS1 activity, differentially regulates HLA-C2-induced secretion of key factors important in vascular remodeling.

**Figure 8.**
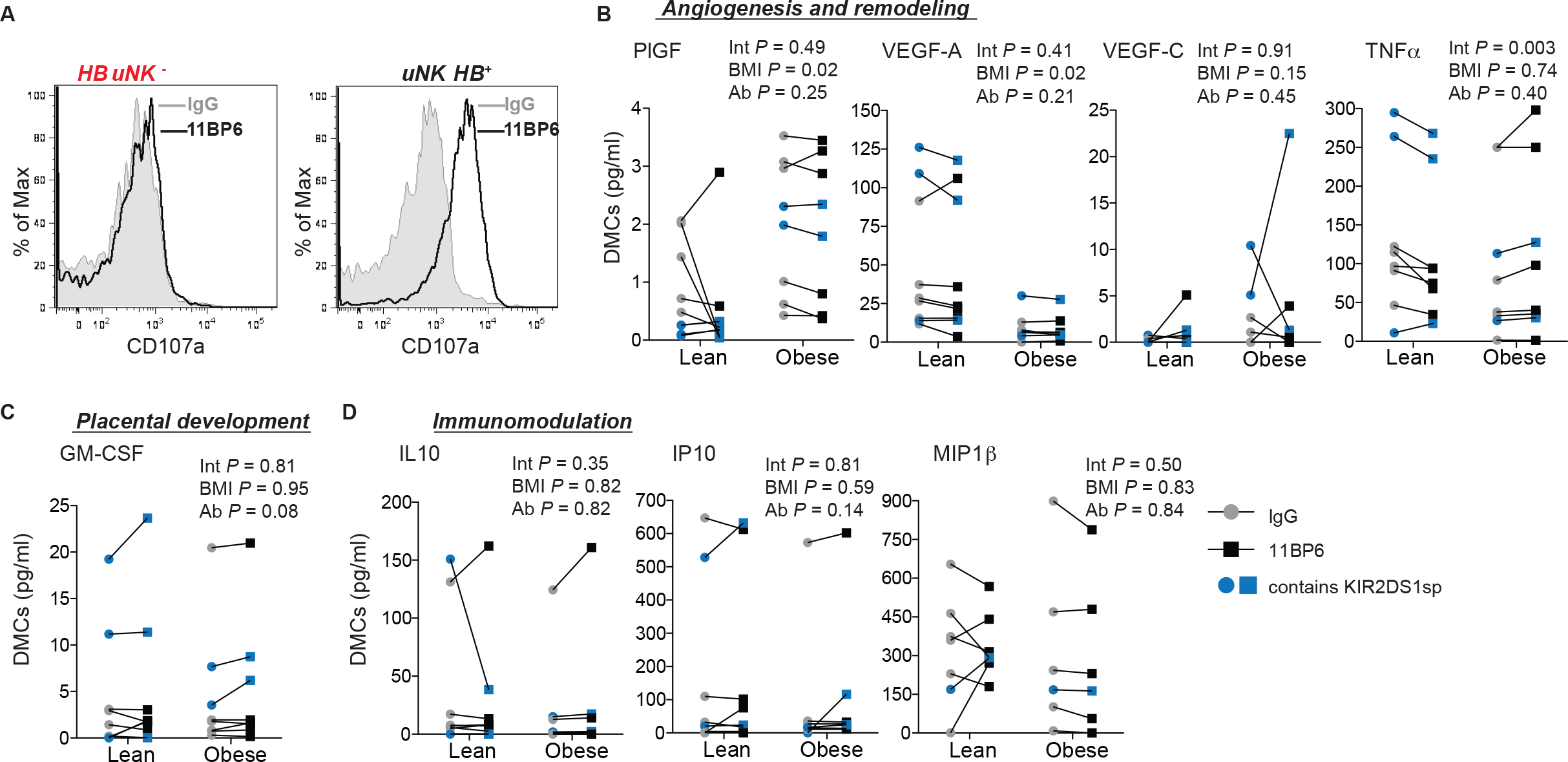
Targeted KIR2DL1/S1 activation differentially controls uNK cytokine production in lean and obese women. **(a)** Representative histograms show CD107a expression in HB^−^ and HB^+^ uNK following 4 hours of KIR2DL1/S1 antibody cross-linking (via clone 11PB6); IgG1 was used a control KIR^+^ cells by flow cytometry. **(b-d)** Scatter plots showing quantification of secreted factors within conditioned media generated from HB^+^ uNK from lean (n = 8) and obese (n = 8) women following KIR2DL1/S1 antibody cross-linking. Secreted factors include **(b)** PlGF, VEGF-A, VEGF-C, TNFα, **(c)** GM-CSF, and **(d)** IL10, IP10, and MIP1β. Significant interactions following crosslinking between uNK from lean and obese subjects were determined via paired repeated measures statistics (two-way ANOVA). Int = Interaction; Ab = 11BP6 Antibody.

## DISCUSSION

In the present study we examined the effects of maternal obesity on uNK activity and function. We show that maternal obesity links with heightened uNK activation/degranulation, and that this change associates with differences in multiple NKRs known to play roles in uNK biology. Specifically, we found that obesity correlates with reductions in NKp46 and NKG2A cell frequencies, while in HB^+^ obese women, exacerbated uNK activity is driven in part by an imbalance in activating KIR2DS1 and inhibitory KIR2DL1 receptor expression. This altered obesity-related KIR2D phenotype, when challenged with ectopic HLA-C2, led to a selective increase in TNFα production and the blunting in inhibition of other key soluble factors. These findings highlight how the condition of obesity is able to instruct functional changes in uNK and establishes insight into how uNK respond and/or adapt to an obesogenic environment.

Our finding that maternal obesity associates with increased uNK activity is both novel and complex. While uNK have a complete arsenal of lytic granules (perforin, granzyme A/B, granulysin), their ability to mount cytotoxic responses against target cells is largely impaired compared to their peripheral NK counterparts^32^. This difference is most likely due to the robust expression of inhibitory NKRs (KIR2DL1, KIR2DL4, NKG2A, LILRB1) in uNK and a hampered ability for uNK to form effective effector-target cell synapses^32^. Inhibitory NKRs facilitate uNK interaction with trophoblast MHC-I antigen (i.e. HLA-C, HLA-E and HLA-G) to generate tolerogenic signals towards the fetus^47^. Even though our work shows that obesity associates with enhanced uNK activity/degranulation and links with altered NKR expression that might seemingly equate to increased cytotoxicity (i.e. decreases in NKG2A & KIR2DL1; increase in KIR2DS1), these changes did not lead to enhanced K562 cell killing. Instead, heightened uNK activity correlated with altered angiokine (increased PlGF, decreased VEGF-A) and KIR2DS1/L1-instructed TNFα production. These findings indicate that although obesity correlates with a shift towards activating NKRs in uNK, inhibitory mechanisms preserving immuno-tolerance/impaired cytotoxicity are largely maintained. Therefore the outcome of increased uNK activity resulting from obesity most likely affects uterine vascular remodeling and angiogenesis. Although our study does not examine this, prior studies showing impaired blood vessel remodeling in rodents subjected to high-fat diets^24,25^ and in obese pregnant women^23,27^ indicate that these vascular defects may be the result of obesity-directed changes in uNK function. Further mechanistic work is therefore required to decipher these interactions.

Our focus on KIR2D(L1/S1) stems from prior research showing the importance of these two receptors in controlling aspects of placental development and in their associations with poor pregnancy outcomes^11,35,48^. Notably, HLA-C2 directed activation of KIR2DS1^49^ and the resulting expression of GM-CSF in 2DS1sp uNK was shown to promote trophoblast invasion^35^. Moreover, a KIR2DS1^+^ containing genotype (i.e. *KIR B* haplotype), in association with fetal HLA-C2 allotype, provides protection from pregnancy disorders like recurrent miscarriage, preeclampsia, and intrauterine growth restriction^36^. In contrast to this, a maternal *KIR AA* genotype (containing 2DL1, but not 2DS1) combined with fetal HLA-C2 increases the risk of aberrant pregnancy outcome^36,48^. Our finding that maternal obesity associates with decreased proportions of 2DL1^+^ uNK and increased expression of 2DS1 in only *KIR B* haplotype (HB^+^) women was somewhat perplexing due to obesity’s predisposition towards having a poor pregnancy outcome^23,50^. Our results indicate that, with respect to KIR2DL1/S1 composition, HB^+^ uNK from obese women may actually be more capable of promoting healthy placentation and may be protective against aberrant pregnancies. It may therefore be that HB^+^ uNK establish an active phenotype as a compensatory mechanism to facilitate successful placentation in obesity. In contrast, HB^−^ uNK (containing 2DL1, but not 2DS1), in the context of obesity, may show aberrant/insufficient activation leading to compromised vascular remodeling and/or inadequate placentation, and thus contribute to or potentiate poor outcomes in these women. Indeed, HB^−^ uNK from obese women were more efficient producers of TNFα following PMA stimulation and secreted lower amounts of GM-CSF compared to HB^−^ uNK from lean women, suggesting these cells may have enhanced cytotoxic potential and/or an impaired ability to promote trophoblast invasion.

Comparison of cytokine/angiokine secretion profiles following KIR2DS1/L1 activation in HB^+^ uNK (2DL1^+^/2DS1^+^) identified TNFα as a cytokine that is differentially regulated with respect to KIR2DL1/S1 balance. Within the context of HB^+^ lean women where 2DL1^+^ uNK proportions are approximately 1.5-fold higher compared to HB^+^ obese women, 2DL1/S1 activation results in decreased TNFα production. In contrast, 2DL1/S1 activation with uNKs from obese women who have fewer 2DL1^+^ cells and also express higher levels of surface 2DS1 leads to an enhancement in, or at the very least, the maintenance of TNFα production. These findings indicate that 2DL1 and 2DS1 play opposing roles in controlling TNFα secretion, a finding that is consistent with previous work showing that 2DS1 engagement with HLA-C2 in CD8^+^ γδT cells induces TNFα^51^. Surprisingly, HLA-C2 interaction did not potentiate an increase in CD107a expression over baseline levels observed in obese uNK exposed HLA-C2-deficient target cells. This inconsistency could be explained by the presence of other activating NKRs responding to target cell exposure (independent of HLA-C2). Moreover, our findings also illustrate how environmental factors, such as obesity, can differentially modulate KIR2D receptor activity following HLA-C interaction. Since factors that regulate KIR2DL1/S1 expression include prior exposure to HLA-C and DNA methylation, and as changes in DNA methylation within promoter/enhancer regions of genes have been associated with obesity^52^, it would be important to examine if obesity regulates KIRs via epigenetic mechanisms.

In addition to KIR2DL1/S1 alterations, our findings also indicate that maternal obesity associates with changes in other non-KIR NKRs that may play roles in modifying uNK function. Specifically, we show that both inhibitory NKG2A and activating NKp46 expression levels and uNK proportions are decreased in obesity. Within the uterine environment, NKG2A’s primary ligand is fetal/maternal derived HLA-E, and this interaction is thought to promote strong tolerogenic/inactivating signals. Thus a loss or reduction in NKG2A signal may be a contributing factor in increased uNK activity in obesity. Our observation of a bimodal distribution of NKp46^+^ uNK in obese women was interesting, and was in stark contrast to the near ubiquitous nature of NKp46 in uNK from lean women. Given that NKp46 is a member of the natural cytotoxicity receptor family and is directly involved in target cell recognition and cytolysis^53^, our finding that NKp46 proportions and expression levels decrease in obesity is contradictory to our finding that obesity also correlates with enhanced surface CD107a/degranulation. However, in pbNK, lower surface levels of NKp46 associate with enhanced activity and exposure to hCMV infection^54,55^. Other studies have shown that increases in NKp46^+^ NK proportions align with enhanced cytotoxicity^56^.

How NKp46 modifications in obese uNK contribute to overall activity and function at this point is unclear. It is tempting to speculate that the combined decrease in NKG2A in uNK may translate to enhanced NKp46 function, a relationship that has been previously identified^56^. Alternatively, loss of the mouse NKp46 orthologue (NCR1) results in penetrant uterine vascular defects defined by impaired angiogenesis, delayed conceptus growth and increased resorption frequencies^57^, findings that are consistent with rodent and human studies showing obesity-related uterine blood vessel defects^27,24,25^. Additionally, NKp46 antibody crosslinking in human uNK induces production of angiogenic VEGF-A and PlGF^30,58^, a finding somewhat consistent with our observation that uNK from obese women secrete less VEGF-A, albeit higher amounts of PlGF. It is possible that this VEGF-A/PlGF imbalance may instruct a shift from PlGF/VEGF-coordinated angiogenesis to aberrant pro-inflammatory signals that are characteristic of conditions associated with of elevated levels of PlGF^59^ and which have been reported to coincide with early pregnancy loss^59^. Therefore it seems possible that uterine vascular defects and altered angiokine profiles seen in obese women may in part be the result of altered NKp46 activity.

uNK are specialized cells that are thought to play essential roles in coordinating and controlling critical processes in pregnancy. While uNK bear some similarities to cytotoxic pbNK counterparts (i.e. contain cytolytic granules), their primary role does not appear to involve the induction of cell-mediated killing. While our study provides evidence that changes within the maternal environment resulting from obesity lead to profound phenotypic and functional differences in uNK, these changes appear to impact predominately angiogenic pathways and do not instruct heightened killing activity. Moving forward, it will be important to examine how the condition of obesity instructs altered uNK function. Identification of obesity-enriched or depleted factors within the maternal-fetal interface that contribute to uNK dysregulation will be important in generating an understanding into the etiology of obesity-related pregnancy disorders.

## METHODS

### Patient recruitment and tissue collection

Informed written consent was obtained from women (19 to 35 years of age) that were undergoing elective pregnancy termination at British Columbia’s Women’s Hospital, Vancouver, Canada. This study was conducted with approval from the Research Ethics Board on the use of human subjects, University of British Columbia (H13-00640). Fresh first trimester decidual tissues (5 to 12 weeks of gestation) and whole blood (N = 184) were collected from participating women having confirmed viable pregnancies indicated by ultrasound-measured fetal heartbeat. Decidual tissue samples were selected based on the presence of a smooth uterine epithelial layer and a course/textured thick spongy underlayer. Patient clinical characteristics i.e. height and weight were additionally obtained to calculate body mass index (BMI: kg/m^2^). Samples were classified as lean (BMI 20-24.9 kg/m^2^; n = 104) or obese (BMI ≥30 kg/m^2^; n = 80). Further information regarding the studied cohorts is depicted in Table 1.

Decidual tissues collected after elective termination of pregnancy where washed extensively in ice-cold phosphate buffered saline (PBS; pH 7.4) after which tissue was finely minced using sterile razor blades and subjected to enzymatic digestion. Decidual leukocyte-enrichment was performed using methods previously described in Perdu *et al*^27^. Briefly, minced decidual homogenates were subjected to 1 h enzymatic digestion at 37 °C in 4 mL of 1:1 DMEM/F12 media (Gibco, Grand Island, NY; 200 mM L-glutamine) containing 1X-collagenase/hyaluronidase (10X stock; StemCell Technologies, Vancouver, Canada), 80 μg/mL DNaseI (Sigma, St. Louis, MO), penicillin/streptomycin, and Anti antimycotic solution (100X dilution). Decidual leukocyte enrichment was obtained via discontinuous Percoll density gradient centrifugation (layered 40%/80%); enriched decidual leukocytes are routinely > 90% CD45^+^ cells as assayed by flow cytometry. Decidual leukocytes were either 1) immediately used to isolate uNK for RNA extraction or short-term cell culture experiments, 2) stored in freezing media (90% FBS + 10% DMSO) to be used for down-stream cell surface marker characterization via flow cytometry, or 3) subjected to cytokine expression or cytotoxicity experiments.

### Flow cytometry analysis

Following washing in fluorescence-activated cell sorting (FACS) buffer (PBS 1X, 1:1000 FVD780, 1% FBS), 1x10^6^ total decidual leukocytes cells were, incubated with FcBlock (Thermo Fisher Scientific, Waltham, MA, USA) for 10 minutes to block unspecific binding. Cells were then incubated with different combinations of fluorescent-conjugated antibodies directed against specific extracellular markers (refer to Supplementary Table 2) for 30 minutes at 4°C.

For *intracellular cytokine detection of TNFα and IFNγ*, cells were stimulated with 10 ng/ml phorbol 12-myristate 13-acetate (PMA) and 500 ng/ml ionomycin (Sigma, St Louis, MO), for 4 hours in the presence of protein transport inhibitors, monensin and brefeldin A (Thermo Fisher Scientific). After stimulation, cells were washed twice with PBS, stained for extracellular markers, fixed and permeabilized with Fixation/Permeabilization buffer (Thermo Fisher Scientific), respectively. Cells were resuspended in 200 μl of PBS and samples were run on a LSRII FACS (BD Biosciences, San Jose, CA, USA). Data was analysed by FlowJo 5.0 (Tree Star, Inc., Ashland, OR, USA).

For the *discrimination of KIR2DS1 and KIR2DL1 cell subsets*, cells were stained for 15 minutes at RT with 10 μl of KIR2DL1. Then, cells were staining with the rest of the markers including KIR2DL1/S1 antibody for 20 minutes at 4°C. Traditional flow cytometry gating strategies for assessing of KIR2DL1/S1 populations gated on CD56^bright^ (total) uNK cells, but in this study we gated on CD56^bright^/KIR^+^ cells. The advantage of our strategy allows recognition of subjects carrying the *KIR2DL3*005* allele, which would be wrongly identified as a KIR2DS1sp subset (Supplementary Figure S8a) due to non-specific binding of monoclonal KIR2DL1/S1 antibodies (clones EB6B and 11PB6)^60^. Analysis of our data using the conventional gating strategy showed no differences (Supplementary Figure S8b) with the results obtained for gated KIR^+^ uNK (Figure 4e).

### Gene expression analysis

Prior to RNA extraction, uNK were enriched using a negative selection exclusion strategy using immuno-labeled magnetic beads cocktails following the manufacturers protocol (EasySep Human NK cell Enrichment Kit; StemCell Technology). Total RNA was prepared from magnetic-bead enriched uNK cells using the RNAqueous Total RNA Isolation Kit following the manufacturer’s instructions (Ambion, Austin, TX). RNA purity was confirmed using a NanoDrop Spectrophotometer (Thermo Fisher Scientific). One microgram of RNA was reverse-transcribed using a first-strand cDNA synthesis kit (QuantaBiosciences Inc., Gaithersburg, MD) and subjected to qPCR (ΔΔCT) analysis, using TaqMan Universal Master Mix II (Life Technologies, Waltham, MA) on an ABI 7500 Sequence Detection System (Life Technologies). Primer sets and Taqman MGB probes (6FAM 5’-3’ MGBNFQ) used for qPCR analysis were as follow: 2DL1F: 5’-*GCAGCACCATGTCGCTCT*-3’; 2DL1R: 5’-*GTCACTGGGAGCTGACAC*-3’; Probe: 5-*ATGCTGACGAACAAGAG*-3’; 2DS1F: 5’-*TCTCCATCAGTCGCATGA*-3’; 2DS1R: 5’-*AGGGCCCAGAGGAAAGTT*-3’; Probe: 5’-*AGGTCTATATGAGAAACC*-3’: 2DS3F; 5’-*TCACTCCCCCCTATCAGTTT*-3’; 2DS3R; 5’-*GCATCTCTAGGTTCCTCCT*-3’; Probe: 5’-*GCCCCACGGTTCT*-3’; 2DS5F; 5’-*GCTCATGGTCATCAGCATGG*-3’; 2DS5R; 5’-*CGACCGATGGAGAAGTTGC*-3’; Probe: 5’-*CACATGAGGGATTCC*. *ACTB* (β-actin) FAM (Hs99999903_m1) from Thermo Fisher. Gene expression was normalized to β-actin levels.

### Genetic typing of KIR2DS1 and HLA-C1/2

Genomic DNA was extracted from peripheral blood mononuclear cells (PBMCs) by DNeasy Blood kit (QIAGEN, Hilden, Germany). PCR amplification was performed using KIR specific sequence primers^48^ to identify the presence or absence of *KIR2DS1* and *KIR2DL1* genes (Supplementary Table 3). HLA-C genes were allotype as either C1 or C2 using sequence specific nucleotides primers as described in^61^ and KIR and HLA-C genotypes were assigned (Supplementary Table 1,3).

### Viral infection of K562 cells with HLA-Cw*0602

Lentiviral vector pHRSIN-HLA-Cw*0602 (kindly donated by Professor John Trowsdale from University of Cambridge, UK) was transiently co-transfected into the HEK293T packaging cell line (kindly provided by Dr. John Priatel, BC Children’s Hospital Research Institute, Canada) using Lipofectamine^®^ 2000 (Thermo Fisher Scientific), together with psPAX2 and pMD2.G vectors (gifted by Dr. Christopher Maxwell, BC Children’s Hospital Research Institute, Canada). Supernatants were harvested 48 hours post-transfection and used to infect K562 cells (provided by Dr. John Priatel, BC Children’s Hospital Research Institute, Canada). Positive clones were sorted based on the expression level of an Emerald green fluorescent protein (eGFP) reporter gene using a BD FACSAria™ IIu cell sorter (BD Biosciences). Cells were sorted directly into growth media.

### Functional assays

An enriched decidual uNK cell fraction was used for all functional assays (> 60% uNK).

*Degranulation/activity* assays were assessed by flow cytometry as described previously^27^. Briefly, 1x10^6^ uNK were cultured alone (non-stimulated), primed with 10 ng/ml PMA, and 500 ng/ml ionomycin, or in co-culture with 2x10^5^ K562 target cells (Effector:Target ratio of 5:1). Next, brefeldin A at 3 μg/ml and monensin at 10 μg/ml from Thermo Fisher were added for the last 3 hours. Uterine NK cells then were stained following the same procedure described in the flow cytometry section. At least 20,000 events were collected in the stopping gate (CD3^−^ FVD780^−^ CD45^+^ CD56^bright^ cells).

*Apoptosis Annexin V/FVD780* assay was performed using Annexin V-PeCy7 antibody (Thermo Fisher Scientific). Culture conditions were the same as described for the K562 target cell degranulation assay (ratio 5:1), without the addition of brefeldin A or monensin. After 4 hours of co-culture, decidual leukocytes and K562 target cells stained for FVD780 and CD45-ef450 for 30 minutes at 4°C. Following washes with Binding Buffer (Thermo Fisher Scientific), cells were stained for Annexin V for 10 minutes at RT, washed in in 200 μl of Binding Buffer 1X, and data was acquired on an LSRII FACS (BD Biosciences); at least 20,000 events were collected.

*Perforin ELISA*. Cell culture supernatants were harvested after 4 hours of decidual leukocyte co-culture with K562 target cells, centrifuged, and snap frozen in liquid nitrogen. Secreted perforin measurements were conducted using Human Perforin ELISA kit (Origene, Rockville, MD, USA) following manufacturer’s instructions. ELISA plates were read using a FLUOstar Optima plate reader (BMG LabTech).

### Multiplex cytokine/angiokine/chemokine array

Decidual leukocytes (DL; > 60% uNK) stimulated by KIR2DL1/S1 receptor ligation using antibody-coated V-bottom 96-well plates (Corning, NY, USA). KIR2DL1/S1 antibody (clone 11PB6; Miltenyi Biotec, Bergisch Gladbach, Germany) or IgG1 (Thermo Fisher) were coated at a concentration of 2.5 μg/ml in 50 mM HEPES buffer at 4°C overnight. Following antibody coating, wells were washed three-times with PBS and 5×10^5^ DL were seeded in RPMI1640 medium containing 10% FBS, 1 mM sodium pyruvate (Life Technologies), 55 nM βME (Sigma), 1% penicillin/streptomycin (Life Technologies), and 1% anti antimycotic (Life Technologies) at 37°C 5% CO_2_. After 4 hours, cell supernatants were collected and analyzed for secreted factors using a custom V-PLEX human biomarkers multiplex assay (PlGF, VEGF-A, VEGF-C, TNFα, GM-CSF, IL10, IP10, and MIP1β) according to manufacturer’s procedures (Meso Scale Discovery, Rockville, MD, USA).

### Statistical analysis

Quantitative PCR (qPCR) gene expression data are presented as mean values ± standard deviation (SD). Flow cytometry and cytokine array data are presented as median values and inter-quartile ranges (IQRs). Differences in continuous variables between two groups were analyzed for statistical significance by non-parametric two-tailed Mann-Whitney U test. Statistical comparisons of uNK cell frequencies and MFIs among (BMI, HLA-C, HB) groups were analyzed using a non-parametric Kruskal-Wallis test; multiple comparisons were controlled for using Dunn’s post test. For identification of BMI-related interactions with KIR2DL1/S1 crosslinking cytokine production, a two-way repeated measures ANOVA was performed. Statistical analyses were performed with GraphPad Prism software (La Jolla, CA, USA).

## AUTHOR CONTRIBUTIONS

AGB designed the research. BC, SP, YM, KC, MM, JA, and AGB, performed experiments and analysed data. BC and AGB wrote the paper. All authors read and approved the manuscript.

## ACKNOWLEDGEMENTS

The authors extend their sincere gratitude to the hard work of staff at British Columbia’s Women’s Hospital’s CARE Program for recruiting participants to our study. We are thankful to Drs. John Priatel (University of British Columbia), Chris Maxwell (University of British Columbia), and John Trowsdale (University of Cambridge) for their generous gifts of the K562 target cell line (Priatel), psPAX2 and pMD2.G packaging vectors (Maxwell), and the pHRSIN-HLA-Cw*0602 lentiviral vector (Trowsdale).

